# Exploring Anti-correlated Resting State BOLD Signals Through Dynamic Functional Connectivity and Whole-brain Computational Modeling

**DOI:** 10.1101/085274

**Authors:** Murat Demirtaş, Matthieu Gilson, John D. Murray, Dina Popovic, Eduard Vieta, Luis Pintor, Vesna Prčkovska, Pablo Villoslada, Gustavo Deco

## Abstract

Resting-state functional magnetic resonance imaging and diffusion weight imaging became a conventional tool to study brain connectivity in healthy and diseased individuals. However, both techniques provide indirect measures of brain connectivity leading to controversies on their interpretation. Among these controversies, interpretation of anti-correlated functional connections and global average signal is a major challenge for the field. In this paper, we used dynamic functional connectivity to calculate the probability of anti-correlations between brain regions. The brain regions forming task-positive and task-negative networks showed high anti-correlation probabilities. The fluctuations in anti-correlation probabilities were significantly correlated with those in global average signal and functional connectivity. We investigated the mechanisms behind these fluctuations using whole-brain computational modeling approach. We found that the underlying effective connectivity and intrinsic noise reflect the static spatiotemporal patterns, whereas the hemodynamic response function is the key factor defining the fluctuations in functional connectivity and anti-correlations. Furthermore, we illustrated the clinical implications of these findings on a group of bipolar disorder patients suffering a depressive relapse (BPD).

## Introduction

The observation of coordinated spontaneous low-frequency fluctuations in motor cortex (Biswal et al. 1995) encouraged interest in the task-free experimental paradigm of fMRI research. Task-free experimental paradigm, known as resting-state fMRI (rs-fMRI), became a useful tool to investigate the alterations in brain connectivity in clinical populations (Greicius 2008; Fox and Greicius 2010; Whitfield-Gabrieli and Ford 2012).

The research on rs-fMRI is mostly based on statistical inferences on the temporal correlations between regional BOLD signals (functional connectivity - FC) or the decomposition of BOLD signals into independent subcomponents (independent component analysis - ICA). Both FC and ICA approaches showed that during resting-state various brain regions co-activate to form the patterns known as resting state networks (RSNs) (Fox and Greicius 2010). In particular, the anti-correlations between two RSNs, namely task-positive (TPN) and task-negative (TNN) networks, gained attention (Fox et al. 2005; Fox and Raichle 2007). The anti-correlated networks provoked methodological concerns such that a common pre-processing procedure, global signal regression, may introduce artefactual anti-correlations (Murphy et al. 2009). Nevertheless, other studies showed these anti-correlated networks using ICA, which does not require global signal regression (Fox et al. 2009; Uddin et al. 2009).

Subsequent studies addressed the temporal variations in functional connectivity (dynamic functional connectivity - dFC) (Chang and Glover 2010; Hutchison et al. 2013; Zalesky et al. 2014; Damaraju et al. 2014; Hansen et al. 2015). In the context of anti-correlations, dFC analyses showed that these correlations waxes and wanes in time (Chang and Glover, 2010).

Various computational models have been proposed to simulate resting-state FC (Cabral, Kringelbach and Deco 2014).These computational models ranged from simple rate models to spiking neural network simulations (Christopher J. Honey et al. 2007; C. J. Honey et al. 2009; Ghosh et al. 2008; Deco et al. 2009; Deco and Jirsa 2012; Messé, Benali and Marrelec 2015). In brief, the computational modeling approach uses structural connectivity (SC) to predict empirically observed rs-FC, therefore it provides a framework that can isolate genuine neuronal activity from the artifacts specific to rs-fMRI.

The structural connectivity between brain regions can be obtained through diffusion weighted imaging (DWI) techniques such as Diffusion Tensor Imaging (DTI) and Diffusion Spectrum Imaging (DSI), among others. Previous studies showed strong relationship between structural connectivity derived from DWI and functional connectivity. However, there are several factors affecting the relationship between SC and FC: First, DWI performs poor at detecting distant
connections and they suffer from gyral bias (Van Essen et al. 2014). Second, DWI does not account for the biophysical properties of the connections such as synaptic conductance. Third, DWI does not provide information on the directionality of the connections, whereas animal studies showed the prevalence of directed connectivity in cortico-cortical connectivity (Markov et al. 2014). All these factors cause poor prediction of the FC at the single subject level, hence they challenge the studies aiming to provide mechanistic understanding of resting state FC.

Effective connectivity (EC) is an estimate for the strengths of directed causal interactions between brain areas and it aggregates the biophysical features and directed connectivity in the brain, thus yielding a richer description of the dynamic interactions of the brain (Karl J. Friston 2011). Several computational models were proposed to infer EC from observed FC (Karl J. Friston 2011; Karl J. Friston et al. 2014; Deco et al. 2014; Gilson et al. 2016).

In this paper, we combined aforementioned techniques to investigate the anti-correlations between brain regions. We tested the hypothesis that the anti-correlated networks may emerge from the time-varying fluctuations in the FC. Using dynamic FC, we quantified the anti-correlation probabilities between brain regions. Then, we used whole-brain computational modeling to investigate the mechanisms behind the anti-correlations. We used recently proposed noise-diffusion model to estimate whole-brain EC and intrinsic node variability from observed FC for each individual subject (Gilson et al. 2016). Furthermore, we studied the effects of hemodynamic response function (HRF) on dFC. Finally, we illustrated the clinical implication of the approach on bipolar disorder patients suffering acute depressive episode.

## Materials and Methods

### Subjects

Patients were recruited from a pool of over 600 patients enrolled in the systematic prospective naturalistic follow-up study of the Bipolar Disorders Program of the Hospital Clinic and University of Barcelona, whose characteristics have been described elsewhere (Popovic et al. 2014). Psychiatric diagnoses were formulated by trained psychiatrists according to DSM-IV-TR criteria and confirmed by Structured Clinical Interview for DSM-III-R-axis I (SCID-I) and axis II-SCID-II (First 2001; First 2002). The severity of the patient's clinical status was evaluated with the 17-item Hamilton Depression Rating Scale (HDRS-17) (Bobes et al. 2003), the Young Mania Rating Scale (YMRS) (Colom et al. 2002) and the Modified Clinical Global Impression Scale for Bipolar Disorder (CGI-BP-M) (Vieta Pascual et al. 2002). Patient’s functioning was rated by the means of Functioning Assessment Short Test (FAST) (Rosa et al. 2007) and medical comorbidities by Charlson Comorbidity Index CGI-BP-M.

We analyzed 13 healthy adults (average age and standard deviation 28.7 ± 3.8 years of age, 10 males) and 8 patients with BP disorder (54 ± 13.2 years of age, 5 males) with a minimum score of 20 on the 17-item Hamilton Rating Scale for Depression (HDRS-17). Patients were included and imaged at the time of suffering an acute depressive relapse (within the first week from onset). Patients were maintained in their previous medication until performing MRI studies. Exclusion criteria included co-morbid Axis I psychiatric conditions, an Axis II diagnosis as determined by the Structured Clinical Interview for DSM-IV Axis II Personality Disorders (SCID-II), a concurrent neurological disorder or an acute medical condition that could interfere with the assessments. The Ethical and Research Committee of the Hospital Clinic approved the study and patients were included after their physicians obtained signed informed consent.

### fMRI acquisition and pre-processing

The subjects were instructed to rest while keeping their eyes closed in a 14-min resting-state scan. Brain images were acquired on a 3 Tesla TrioTim scanner (Siemens, Erlangen, Germany) using the 8-channel phased-array head coil supplied by the vendor. A custom-built head holder was used to prevent head movement, and earplugs were used to attenuate scanner noise. High-resolution three-dimensional T1-weighted magnetization prepared rapid acquisition gradient echo (MPRAGE) images were acquired for anatomic reference (TR=2200ms, TE=3ms, FA=7°, 1.0mm isotropic voxels). T2-weighted scan was used in order to identify pathological findings (TR=3780ms, TE=96ms, FA=120°, voxel size 0.8×0.6×3.0mm, 3.0mm thick, 0.3mm gap between slices, 40 axial slices). Functional data were acquired using a gradient-echo echo-planar pulse sequence sensitive to blood oxygenation level-dependent (BOLD) contrast (TR=2000ms, TE=30ms, FA=85°, 3.0mm isotropic voxels, 3.0mm thick, no gap between slices). The spatial resolutions of the scans were 2.0×2.0×3.0 for the bipolar disorder patients and 2.5×2.5×2.5 for the healthy control subjects. Both groups were resampled at 2.0×2.0×2.0 spatial resolution for the preprocessing. fMRI data were preprocessed using SPM8 (http://www.fil.ion.ucl.ac.uk/spm) including compensation of systematic, slice-dependent time shifts, motion correction, and normalization to the atlas space of the Montreal Neurological Institute (MNI) (SPM8). We used the SPM connectivity toolbox *Conn* (http://web.mit.edu/swg/software.htm) with a temporal filtering that remove constant offsets and linear trends over each run but retained frequencies below 0.1 Hz. Data was spatially smoothed using a standard 8 mm full-width half-maximum Gaussian blur. The brains were parceled into 82 regions based on scale 33 Lausanne Atlas.

### Acquisition of Structural Connectivity Matrices

DSI data was acquired in the same scanning session with 515 gradient directions at a max b-value of 8000 s/mm2 (TR/TE=8200/164 ms) and voxel size of 2×2×3mm^3^ maximum diffusion gradient intensity: 80 mT/m with a total acquisition time of 35.42 min. The details were described elsewhere (Prčkovska et al. 2016).

### Dynamic Functional Connectivity and Anti-correlation Probabilities

We used sliding window analysis approach to construct dynamic functional connectivity tensors (dFC). We used a sliding window of size 30TR (60 seconds) with 5TR (10 seconds) step size. For each window FC was estimated using Pearson’s correlation coefficient. For the simulated time-series, we used a sliding window size of 300TR (keeping the 5TR step size). This was done due to computational efficiency because longer simulation time was needed for the correlation coefficient between simulated signals to stabilize.

For both empirical and simulated time series the histograms of correlations coefficients were estimated for each functional connectivity. Then we quantified the anti-correlation probabilities (ACP) as the ratio of observing correlation coefficients lower than a negative threshold (in the paper we used r = -0.25) across time for all subjects. To summarize the overall ACPs per subject, we used the average across all ACPs.

Global average signal (GAS) was calculated as the mean across BOLD signals of all regions at each time point. The GAS spatial patterns were estimated as the correlation coefficient between BOLD signal of each region with GAS.

### Computational Model

We used the computational model based on linear noise diffusion to estimate the directed effective connectivity (EC) from the observed data, which was previously implemented on resting-state fMRI signals (Gilson et al. 2016). The model described the time courses of rs-fMRI BOLD signals as a linear noise diffusion process. The equations comprised following coupled differential equations:

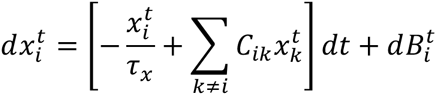

where 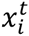 denotes the activity in each brain region with a decay time constant *τ*_*x*_, the activity in each node is coupled by the connectivity matrix, *C*, and 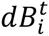 denotes a Wiener process with factor *σ*_*i*_.

When the system has a stable fixed point, the self-consistency equations for the mean 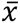 and the covariance *Q*^*τ*^ can be derived as follows:

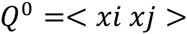

Where < > denotes averaging over randomness. The Jacobian of the system, *J*, was calculated using:

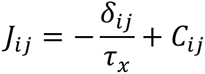

Here *δ*_*ij*_ denotes Kronecker delta, *δ*_*ij*_ = 1, if i = j, 0 otherwise.

For zero time shift, we obtained the Lyapunov equation for the steady-state of the second-order fluctuations using the equation:

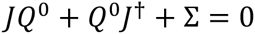

Given the noise matrix ∑, with the diagonal terms of 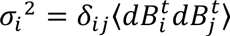, zero time shift covariance matrix, *Q*^0^, can be obtained by solving the equation. Dagger symbol denotes the matrix transpose. Similarly the covariance with time shift *τ* (here we used *τ* =1) can be calculated as:

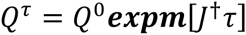

This framework allowed estimation of directed effective connectivity using Lyapunov optimization, which is described as follows:

Given the observed covariance matrices (objective matrices) 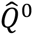 and 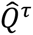, we aimed to optimize the Lyapunov function, V, for C:

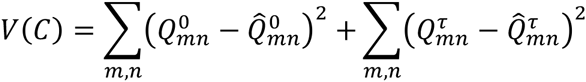

The Lyapunov function is positive definitive and it becomes zero only if both covariance matrices are equal to the objective values. We started with zero connectivity matrix and at each step updated it, *C_ij_*, such that the model covariance approaches to the objective counterparts. Therefore the desired change for the model covariance can be defines as:

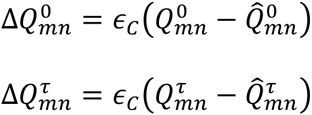

where *∈*_*C*_ is an arbitrary small positive number. The corresponding update for the Jabobian can be derived giving the following equation:

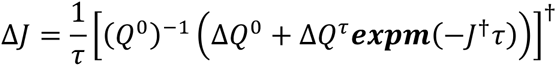

Finally, the connectivity matrix was updated for only existing connections (*C*_*ij*_ > 0) using:

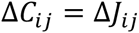

To estimate the node variances, at each optimization step the noise matrix, ∑, is estimated by using the equation:

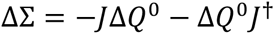

To avoid overfitting of the model, we constraint the model to non-zero connections in the observed SC matrices averaged across subjects.Furthermore, the effective connectivity was bounded between 0 and 1. Prior to the optimization procedure decay time constants, *τ*_*x*_, were estimated by fitting to the autocovariances of the empirical BOLD time series.

### BOLD Transformation of Simulated Time-Series

The activity computed via the generative model can be transformed into a BOLD-fMRI signal using the Balloon-Windkessel hemodynamic model (Buxton, Wong, and Frank 1998; K. J. Friston, Harrison, and Penny 2003). According to this model, the vasodilatory signal *s_i_*, in node *i* increases due to the neuronal activity *z_i_*. Thus the blood volume *v_i_*, deoxyhemoglobin content *q_i_* and inflow *f_i_* change in response to the vasodilatory signal. The set of coupled differential equations is shown below:

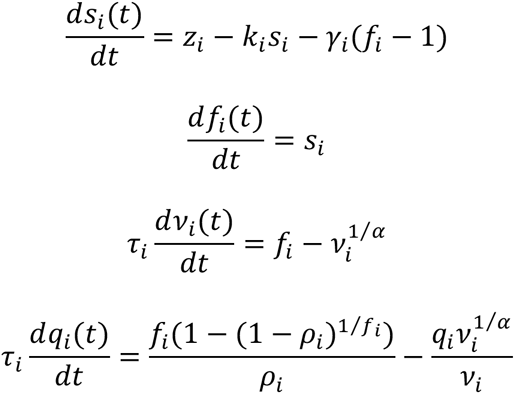

Finally, the BOLD signal yi is computed as a static nonlinear function of blood volume and deoxyhemoglobin content:

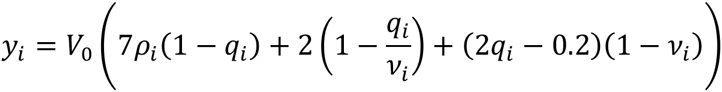

where *ρ* is the resting oxygen extraction fraction and *V*_*0*_ = *0.02* is the resting blood volume fraction.

### Statistical Analyses

The group comparisons for average FC, average ACP, SC strength, FC strength, EC output and input strengths, and nodal variance were done using a permutation t-test with 10000 permutations. Age of subjects was regressed out from each parameter before performing permutation tests. The p-values of the connectivity strength comparisons were corrected for multiple comparisons using false discovery rate (FDR) approach (Benjamini and Yekutieli 2001).

The associations between average ACP and FC strength, EC output and input strengths and nodal variance were estimated using partial correlations controlled for average FC. The correlations were controlled for average FC to isolate the effect of ACP considering the high negative correlation between ACP and FC.

## Results

### Functional connectivity distributions and anti-correlation probability

We investigated the relationship between dynamic FC and anti-correlated networks using sliding window analysis approach. The grand-average FC, without global signal regression, did not exhibit strong negative correlations between brain regions (figure 1a). Given the distributions of the correlation coefficients between brain regions across time (figure 1b), we introduced an alternative measure for anti-correlations (anti-correlation probabilities; ACP) that quantifies the probability of the correlation coefficients being lower than the threshold (r < -0.25) (figure 1d). We found significant negative correlation between average ACP and FC (r = -0.91, p < 0.001)(figure 1c,f). The cumulative average of ACPs decreased monotonously and it stabilized suggesting that ACPs were time dependent (figure 1e). Furthermore, we investigated the relationship between global average signal (GAS) fluctuations and ACPs. We found significant negative correlation between GAS variance and average ACP across subjects (r = -0.6, p-value= 0.03). At each sliding window, minimum GAS was significantly correlated with average ACP (r = 0.46, p-value <0.001), whereas maximum GAS was significantly anti-correlated with average ACP (r = -0.53, pvalue< 0.001). Moreover, to justify further analysis, we checked the relationship between GAS spatial patterns and SC strength and we found significant correlation (r = 0.3, p-value = 0.007).

**Figure 1.**
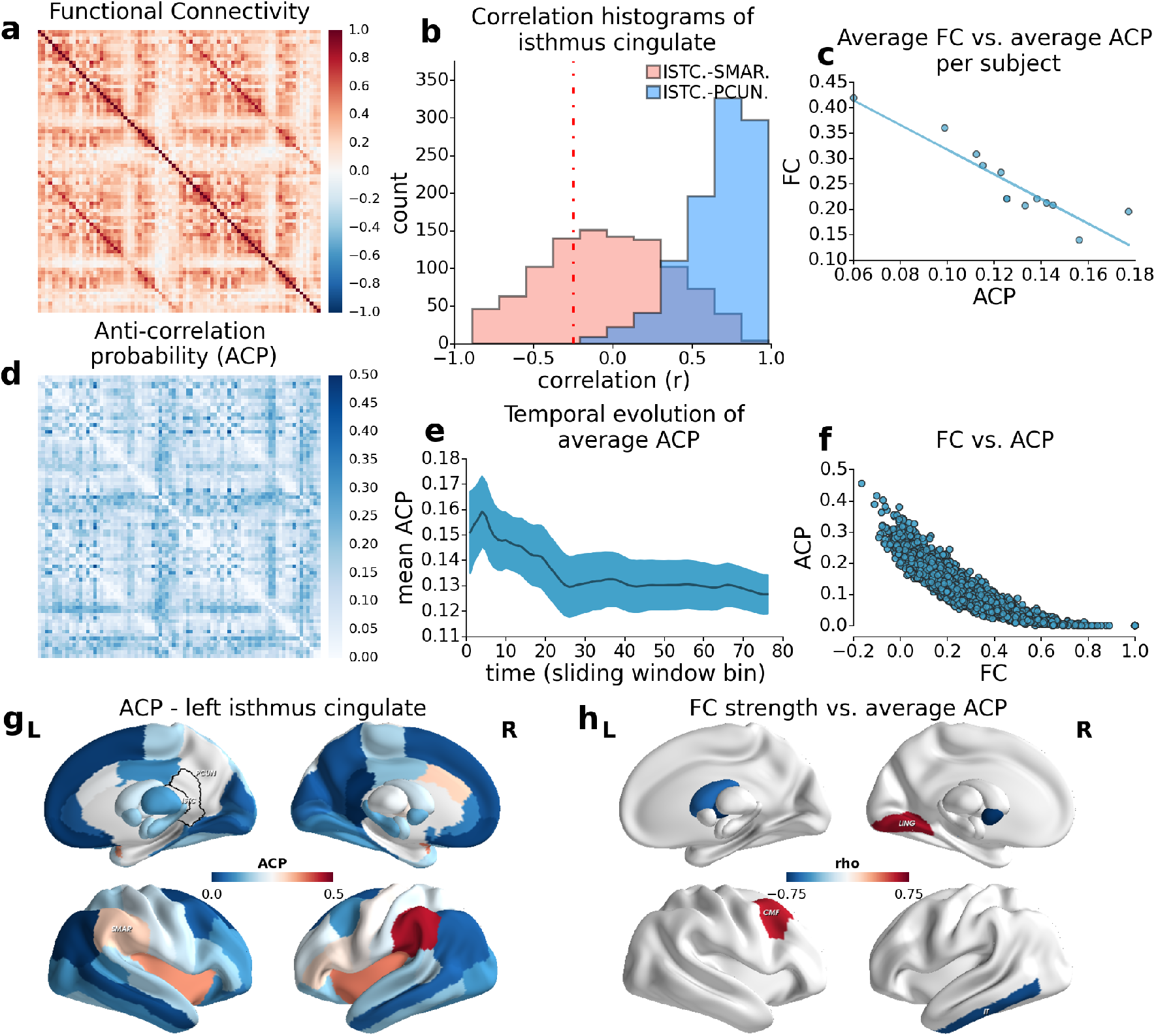
FC distributions and anti-correlation probabilities (ACP). A. Grand-average Functional Connectivity. B. Histograms for FC between left isthmus cingulate and left supramarginal gyrus (salmon), left isthmus cingulate and left precuneus (blue). Red dashed line indicates the threshold. C. The relationship between average FC and average ACP per subject. C. Average ACPs for each connection pair. E. Cumulative mean ACP with respect to sliding window time bin (shaded region represents standard deviation). F. The relationship between FC and ACP per connection pair. G. ACPs of left isthmus cingulate seed (bordered region). Hot colors indicate higher ACP. H. Partial correlation coefficients between FC strengths of each node versus average ACP (controlling for average FC).

We checked qualitatively whether the ACPs showed any patterns similar to task-positive (TPN)and task-negative (TNN) networks using left isthmus cingulate as seed-region (figure 1g) (see supplementary table 1 for details). ACP of left isthmus cingulate was the lowest with right isthmus cingulate, bilateral precuneus, medial orbiral frontal cortex, superior frontal gyrus, inferior parietal cortex. In contrast, left isthmus cingulate seed exhibited the high ACP with bilateral supramarginal gyrus, insula, caudal anterior cingulate cortex, right parstriangularis, and right pars opercularis. Low and high ACP patterns suggested transient emergence of TPN and TNN in dFC.

We investigated the relationship between FC strength of each node and ACPs (figure 1h). The correlations were controlled for the average FC of each subject due to high correlation between average FC and average ACP. FC strengths of right lingual gyrus (rho = 0.64, p-value = 0.02) and left caudal middle frontal gyrus (rho = 0.59, p-value = 0.03) was positively correlated with ACP. FC strengths of left pallidum (rho = -0.86, p-value < 0.001), right accumbens (rho = -0.74, p-value < 0.001), right inferior temporal gyrus (rho = -0.74, p-value = 0.01),right putamen (rho = -0.67, p-value = 0.01) and left caudate (rho = -0.60, p-value = 0.03) were negatively correlated with ACP (see supplementary table 2 for details).

### Computational modeling of anti-correlation probability

We estimated the effective connectivity and node variance using a linear noise-diffusion model. The average correlation coefficient between empirical and simulated values was r = 0.54 (std = 0.083) for FC, and r = 0.31 (std = 0.073) for ACP (figure 2a-c). However, the simulated dynamic FC exhibited narrow FC distribution, which did not correspond to the empirical observations (figure 2d). The simulated ACPs of left isthmus cingulate showed the lowest peak value with precuneus and the highest peak value with lateral orbital frontal cortex. Therefore, despite the similarities in the empirical and simulated patterns, the model did not showed negative correlations.

**Figure 2.**
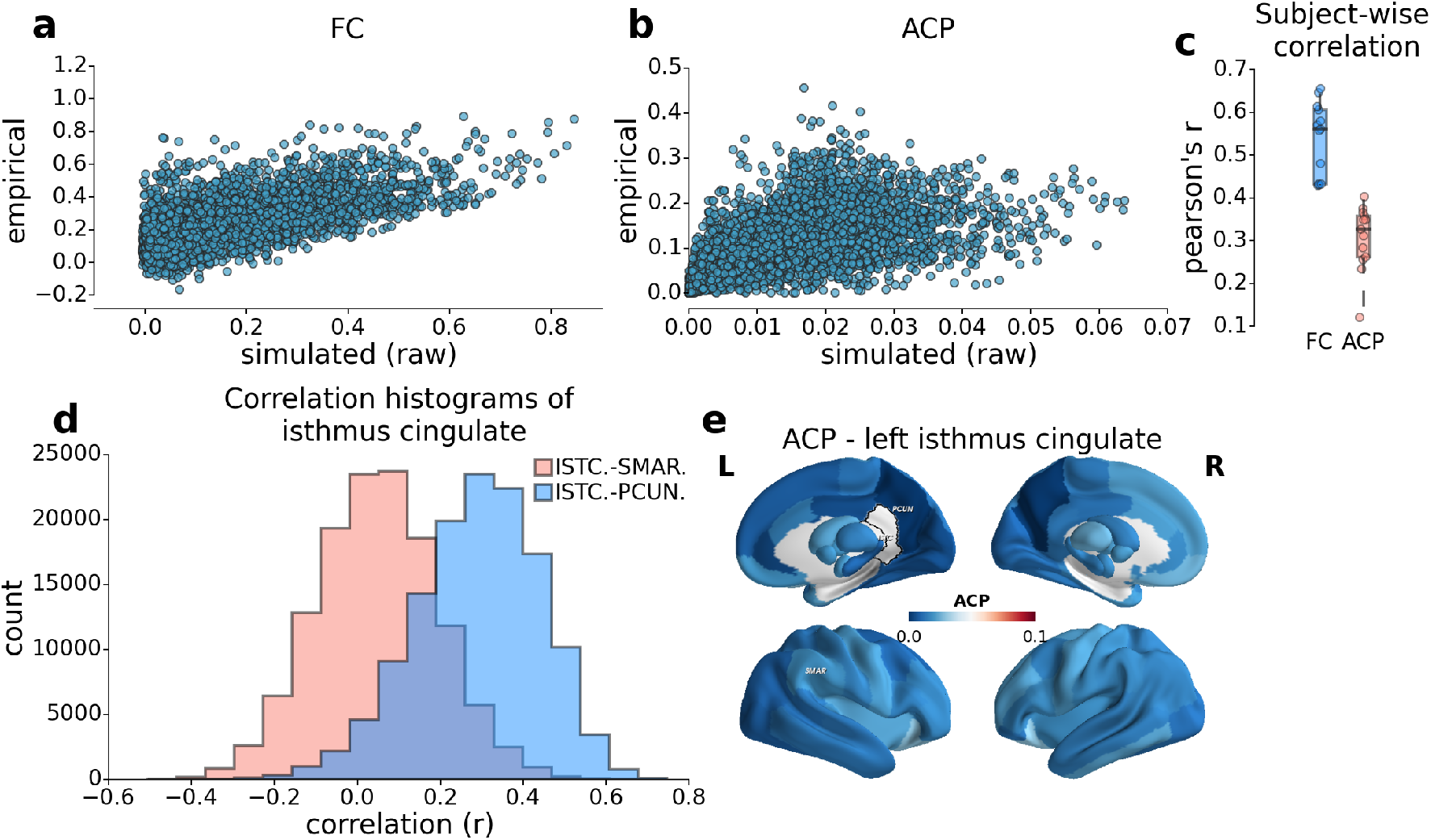
Simulated measures based on inferred EC for all subjects. The relationships between empirical and simulated FC (A), ACP (B). C. Pearson’s correlation coefficients between empirical and simulated FC and ACP per subject. D. Histograms for simulated FC between left isthmus cingulate and left supramarginal gyrus (salmon), left isthmus cingulate and left precuneus (blue) (simulated counterpart of figure 1B). E. ACPs of left isthmus cingulate seed (simulated counterpart of figure 1G).

We also investigated the role of the hemodynamic BOLD responses on FC dynamics (figure 3). BOLD transformed simulations exhibited similar correspondence between empirical and simulated FCs (r = 0.56, std = 0.097) with no significant differences between simulations (T-statistic = -0.71, p = 0.48). Nevertheless, the similarity between empirical and simulated ACP increased significantly (r = 0.39, std = 0.092) (T-statistic = -2.26, p = 0.03). Furthermore, the FC distributions of the BOLD transformed model were qualitatively similar to the observed data. After BOLD transformation, precuneus remained as the region which showed the highest ACP with left isthmus cingulate. The highest simulated left isthmus cingulate ACP was observed with entorhinal cortex.

**Figure 3.**
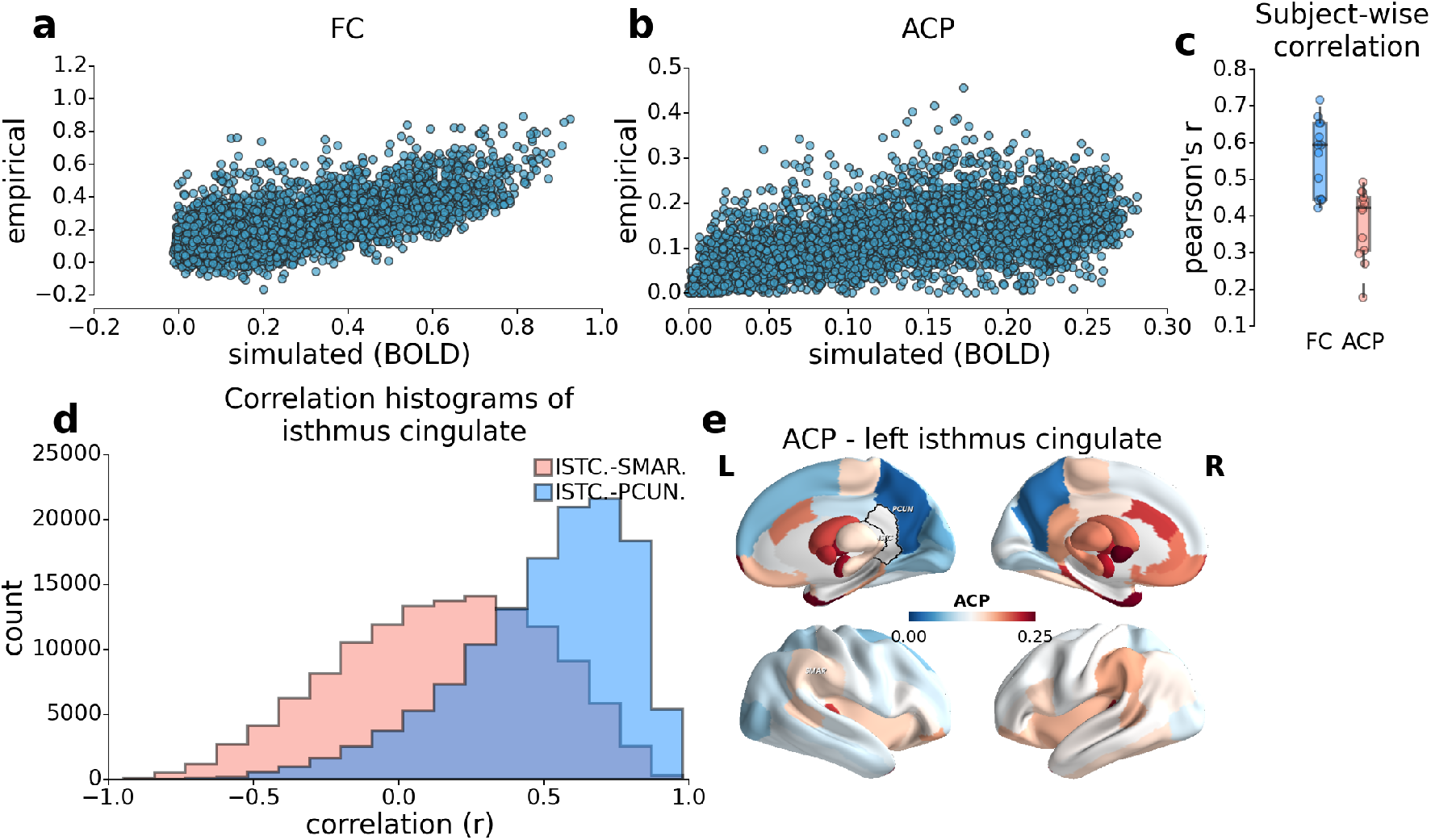
Simulated measures based on inferred EC for all subjects after transforming the signal to BOLD (see figure 2).

### Effective connectivity and anti-correlation probability

We studied the relationship between empirical ACPs and estimated model parameters (figure 4)(see supplementary table 2 for details). Controlling for average FC, we computed the partial correlations between average ACP and EC output strength, EC input strength, and node variance. Average ACP was positively correlated with EC output strength of right paracentral gyrus, and negatively correlated with left pallidum, right precentral gyrus, medial orbital frontal cortex, para-hippocampal gyrus, inferior temporal gyrus, amygdala, and accumbens (figure 4).

**Figure 4.**
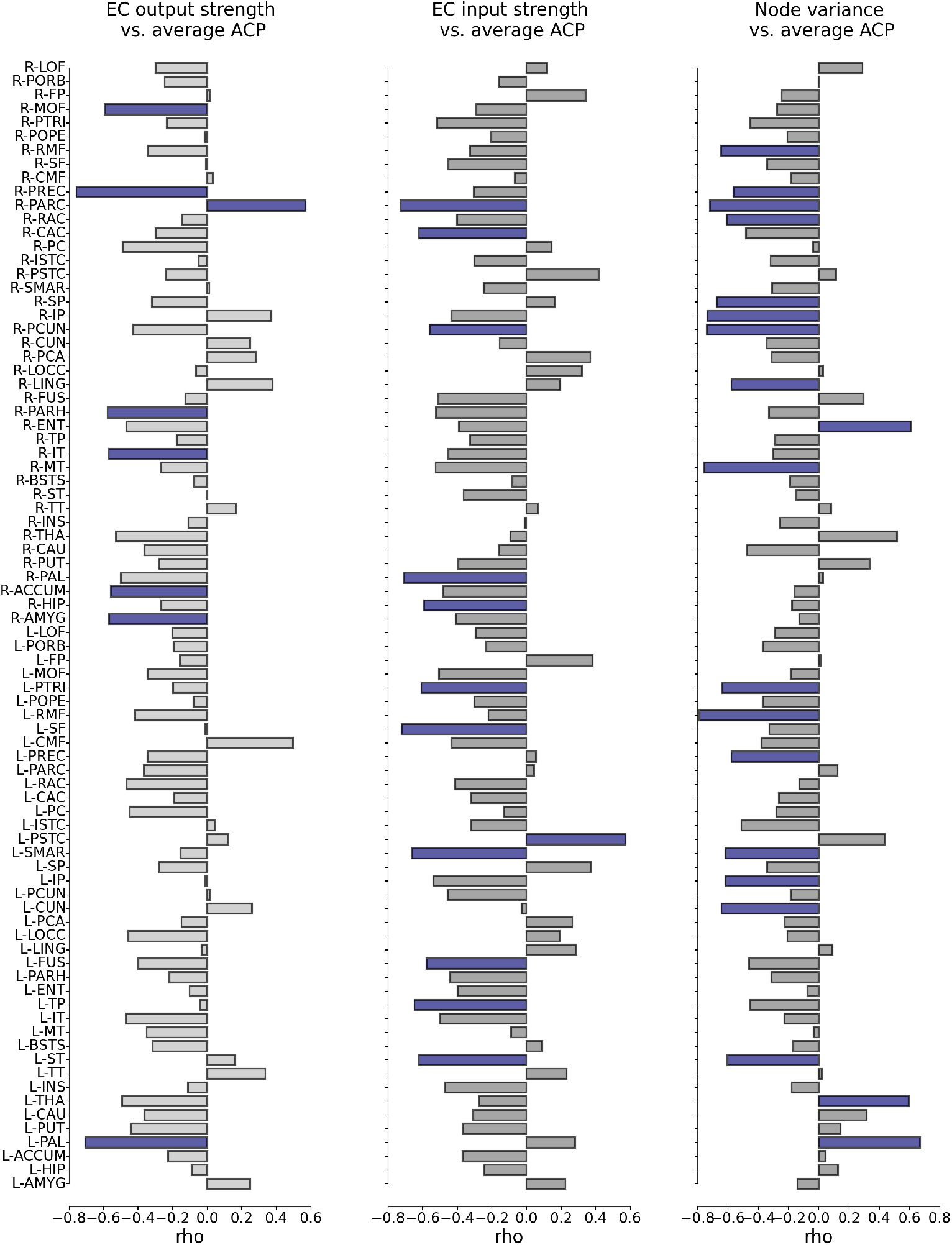
Partial correlation coefficients between average ACP versus EC output strength (first column), EC input strength (second column), and node variance (third column) controlling for average FC.

EC input strength was positively correlated with average ACP in left postcentral gyrus, and negatively correlated with average ACP in right paracentral gyrus, left superior frontal gyrus, right pallidum, left supramarginal gyrus, left temporal pole, left superior temporal gyrus, right caudal anterior cingulate cortex, left pars triangularis, right hippocampus, left fusiform gyrus, and right precuneus (figure 4). In summary, EC output strength from sensory motor regions modulated ACPs, whereas EC input strength to default mode and frontal/parietal networks modulated ACPs. In addition, among subcortical regions negative correlations with ACP was significant for output strengths of amygdala and accumbens, whereas it was significant for input strengths of hippocampus and pallidum.

The correlations between average ACP and intrinsic node variability was positive in right pallidum, right entorhinal cortex, left thalamus. Negative correlations were observed in rostral middle frontal, inferior parietal, precentral gyrus, right medial temporal gyrus, right precuneus, right paracenral gyrus, right superior parietal, left cuneus, left pars triangularis, left supramarginal gyrus, right rostral anterior cingulate, left superior temporal gyrus, and left lingual gyrus (figure 4). Overall, intrinsic node variability was more influential on ACP than EC.

### Clinical Application of the Model

To investigate the clinical relevance of the model, we compared the empirical (FC strength, SC strength, average FC, average ACP), and simulated (EC output strength, EC input strength, node variance) measures of bipolar disorder patients during the depressive relapse and healthy controls. We found no significant differences in average FC between groups (T-statistic = 1.84, p-value = 0.08, permutation t-test, 10000 permutations), while the average ACP was significantly increased in BDP group (T-statistic = -3.72, p-value < 0.01, permutation t-test, 10000 permutations)(figure 5). In BPD group, we found significant decreases in SC of left posterior and isthmus cingulate, and in FC of right precentral gyrus, medial and transverse temporal gyri, left pars-orbitalis, precuneus, precentral gyrus, bank of superior temporal and superior temporal gyri (see supplementary table 3 for details). None of these alterations survived the correction for multiple comparisons.

We found no significant differences in EC output and input strengths. In contrast,
intrinsic node variances were significantly increased in BPD group (see supplementary table 3 for details). However, similar to the differences in SC and FC, none of the results survived the correction for multiple comparisons.

**Figure 5.**
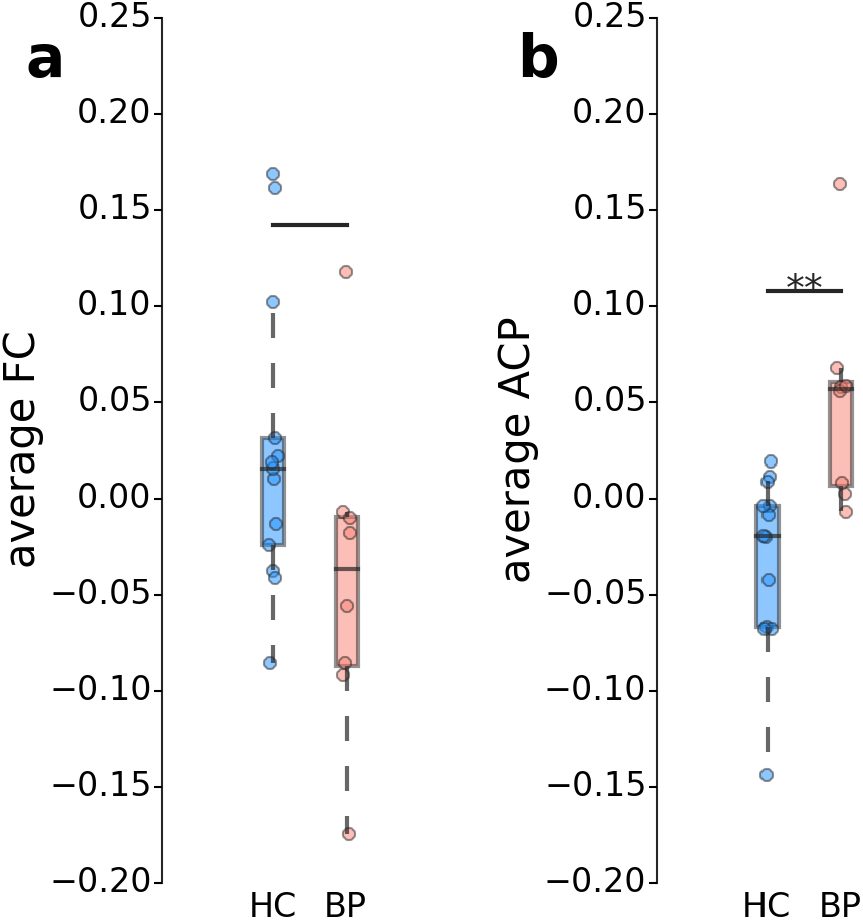
Statistical comparisons between healthy controls and BPD patients for average FC (A) and average ACP (B).

## Discussion

Anti-correlations between brain regions during spontaneous activity have been a controversial topic in resting-state fMRI research. Global signal regression (GSR) was claimed to be responsible for the anti-correlations that are observed in rs-FC. The intuition behind this claim was that GSR artificially introduces anticorrelations by shifting the grand-average FC distributions around zero (Murphy et al. 2009). Several studies showed the presence of anti-correlated networks independent component analysis (ICA) approach, which does not require GSR (Fox et al. 2009). Other studies that use dynamic FC approach suggested that the anti-correlations are not stationary and they emerge transiently. In parallel, clinical rs-fMRI studies showed that GSR qualitatively alters the results in clinical populations (Yang et al. 2014), and that GAS fluctuations play key role in dynamic FC (Demirtaş et al. 2016). In contrast, a recent paper showed that GAS fluctuations substantially reflect variability in the physiological signals such as respiration and hearth rate (Power et al. 2016).

In this paper we combined dynamic functional connectivity and whole-brain computational modeling approaches to reconcile the controversy around anticorrelations. Using dynamic FC we computed anti-correlation probabilities (ACPs) between brain regions. ACPs exhibited a strong negative correlation with grand-average FC. This was an expected result because weakly correlated regions would become anti-correlated more likely (and vice versa) due to the observation noise on FC across time. The results suggested that ACPs reflect patterns that are reminiscent of task-negative (TNN) and task-positive (TPN) networks. The regions forming these two networks exhibited negative correlations up to 45% of time. Previous studies already showed that the anticorrelated networks may wax and wane across time. Adding to these findings, we showed that the anti-correlations more likely to occur at low-GAS states than they do at high-GAS states and that they are substantially related to variability of GAS. These results links the anti-correlations introduced by GSR and the GAS itself. We predict that even in ICA approach the observed anti-correlations are related to GAS fluctuations.

One important implication of these findings is that anti-correlated RSNs might merely reflect the observation noise in FC distributions even without GSR. We investigated this hypothesis using a whole-brain computational model based on linear noise diffusion. The model allowed us to study the role of effective connectivity (EC) and nodal dynamics in generating observed grand average FC for each subject. Despite the high similarity between empirical and simulated FC and ACP, the linear model did not reflect the observed FC distributions. Being specific, in the simulated time-series the patterns of ACPs were relatively preserved but the variability of FC was lower than the empirical data. The negative correlations between regions in simulated time-series reached up to 3%, which was much lower that the observed values. For this reason, we studied the role of the nonlinearities of the hemodynamic response function (HRF). The similarity between empirical and simulated BOLD signals did not changed significantly after BOLD transformation. However, it dramatically increased the variability of FC in time leading to similar FC distributions and ACPs (up to 25%) to the observed values. Furthermore, the correlation between empirical and simulated ACP was significantly higher after the BOLD transformation. The results support some recent studies that showed relationship between dynamic FC and physiological signals (Nikolaou et al. 2016), and between negative BOLD response and anti-correlations (Bright et al. 2014). Furthermore, these results are consistent with the findings suggesting strong relationship between GAS fluctuations and physiological signal (Power et al. 2016). Nevertheless, our findings suggest that the relationship between GAS and physiological signal does not necessarily imply GAS fluctuations reflect non-neuronal artifacts. Indeed, we also showed significant correlation between SC and GAS correlation maps. Based on these results we speculate that in addition to genuine GAS fluctuations of neuronal origin, the underlying neuronal signal may be enhanced/attenuated due to the fluctuations in hemodynamic response function of physiological origin. The partial correlations between average ACP and the model parameters suggested the role of subcortical and sensory-motor regions on anti-correlations. This observation is consistent with previous findings on the role of subcortical regions as a hub for anti-correlations (Gopinath et al. 2015). The analyses also suggested higher anti-correlation power associated with lower EC output strength from sensory-motor regions and lower EC input strength to higher association regions. Furthermore, the influence of intrinsic node variance on ACPs were more widespread than EC. In particular, node variance of thalamus and pallidum showed positive correlations with ACP, while frontal and parietal regions showed negative correlations. These finding suggests regulatory role of cortical-subcortical interactions across the cortical hierarchy on anti-correlation power.

We showed that observation noise over FC due to the nonlinearities in HRF is a factor contributing to the dynamic fluctuations in FC and anti-correlated regions in the brain. Nevertheless, the model fit for ACP was weaker than the grand average FC. Moreover, the model performed poor on estimating the correlations/anti-correlations in subcortical regions. The propose framework does not allow to investigate whether this is due to the non-homogeneities in HRF or peculiarities of subcortical structures such as inhibitory connections, oscillations or subdivisions.

Finally, we demonstrated the clinical relevance of the model on BPD patients with depressive relapse. After regressing out subject age as a confounding variable and correcting the p-values for multiple comparisons, we found no significant differences between groups (for average FC, SC strength, FC strength, EC-input and -output strengths, node variance). However, average ACP remained significantly higher in patients with BPD. In light of the previous discussion, we speculate that abnormal HRF in BPD subjects may play role in the alterations observed in resting-state dynamic FC, which might be applied to many other disorders.

This study has various limitations and considerations. First, the sample sizes are not large enough to generalize the results, especially for the clinical analysis. The main reason for small sample size is that bipolar disorder with depressive relapse is a specific case, hence it is also an unexplored clinical population. Second, the study does not involve a quantitative description of the resting state networks as the computational approach relies on the simulation of parcellated brain regions as nodes. Third, the model-based inference of the whole-brain EC has limitations due to the high number of parameters. The face validity of the optimization procedure was illustrated previously in a larger sample (Gilson et al. 2016), and we further addressed the problem of overfitting by constraining the optimization to positive, non-zero connections in anatomical connectivity.

---

**Supplementary Materials.**
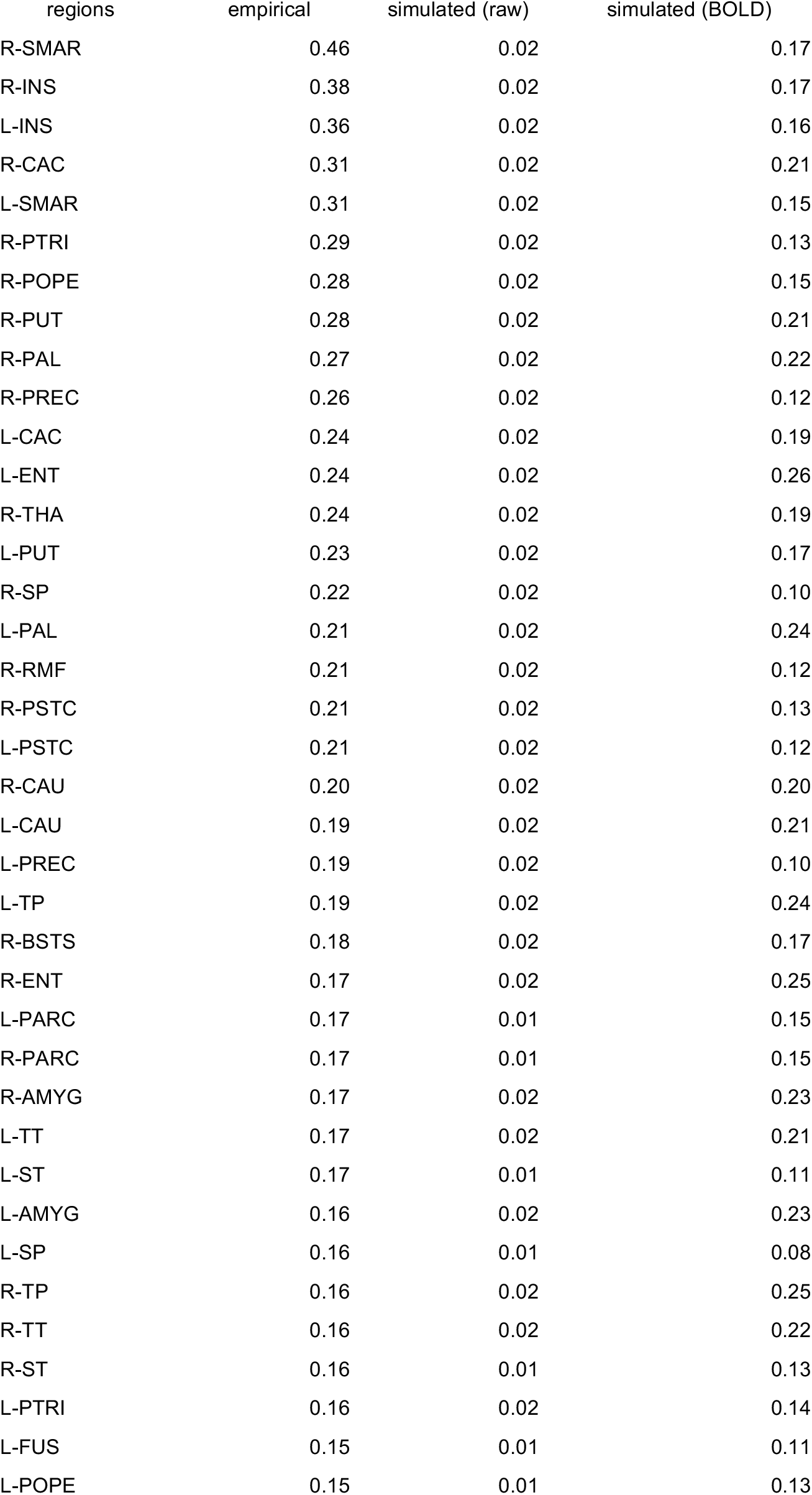
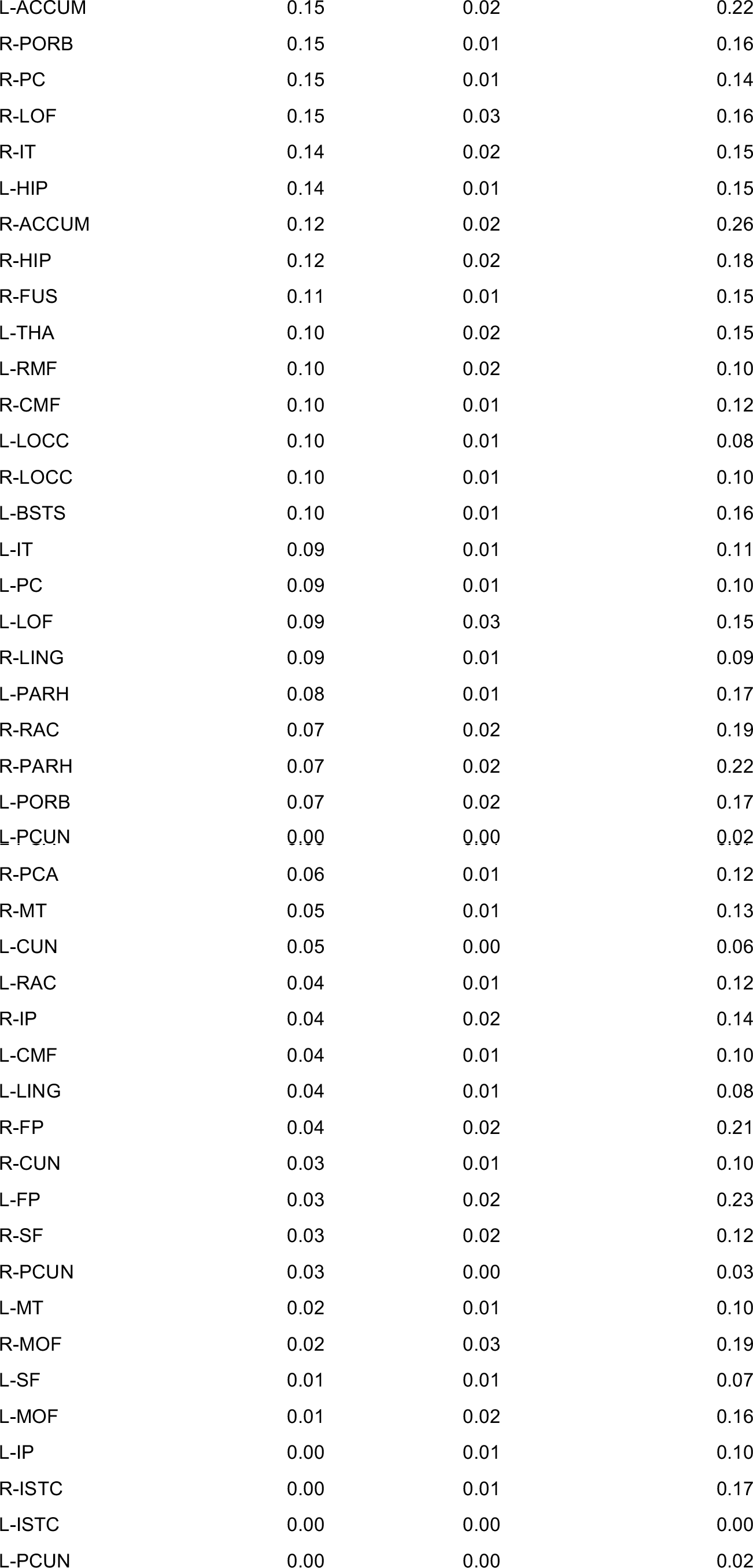
Supplementary Table 1 - ACP of left isthmus cingulate (ordered)

Supplementary Table 2 - Partial correlations between average ACP controlled for average FC.

| regions | Functional Connectivity |  | EC input-strength |  | EC output-strength |  | Node variance |  |
| --- | --- | --- | --- | --- | --- | --- | --- | --- |
|  | rho | p-values | rho | p-values | rho | p-values | rho | p-values |
| R-LOF | 0.09 | 0.78 | 0.12 | 0.70 | -0.30 | 0.32 | 0.29 | 0.34 |
| R-PORB | 0.12 | 0.69 | -0.16 | 0.60 | -0.25 | 0.42 | 0.00 | 0.99 |
| R-FP | 0.09 | 0.78 | 0.34 | 0.25 | 0.02 | 0.95 | -0.25 | 0.42 |
| R-MOF | -0.26 | 0.40 | -0.29 | 0.34 | -0.59 | 0.03 | -0.28 | 0.36 |
| R-PTRI | 0.30 | 0.33 | -0.52 | 0.07 | -0.23 | 0.44 | -0.45 | 0.12 |
| R-POPE | 0.17 | 0.58 | -0.20 | 0.51 | -0.01 | 0.96 | -0.21 | 0.49 |
| R-RMF | 0.04 | 0.90 | -0.33 | 0.28 | -0.34 | 0.25 | -0.65 | 0.02 |
| R-SF | 0.43 | 0.14 | -0.45 | 0.12 | -0.01 | 0.98 | -0.34 | 0.25 |
| R-CMF | 0.33 | 0.28 | -0.07 | 0.82 | 0.03 | 0.92 | -0.18 | 0.55 |
| R-PREC | 0.36 | 0.23 | -0.30 | 0.31 | -0.76 | 0.00 | -0.56 | 0.05 |
| R-PARC | 0.48 | 0.10 | -0.73 | 0.00 | 0.57 | 0.04 | -0.72 | 0.01 |
| R-RAC | -0.11 | 0.72 | -0.40 | 0.17 | -0.15 | 0.63 | -0.61 | 0.03 |
| R-CAC | -0.44 | 0.14 | -0.62 | 0.02 | -0.30 | 0.32 | -0.48 | 0.10 |
| R-PC | -0.27 | 0.38 | 0.15 | 0.64 | -0.49 | 0.09 | -0.04 | 0.90 |
| R-ISTC | -0.10 | 0.74 | -0.30 | 0.31 | -0.05 | 0.87 | -0.32 | 0.29 |
| R-PSTC | 0.45 | 0.12 | 0.42 | 0.16 | -0.24 | 0.43 | 0.11 | 0.72 |
| R-SMAR | 0.23 | 0.46 | -0.25 | 0.41 | 0.01 | 0.96 | -0.31 | 0.31 |
| R-SP | 0.48 | 0.10 | 0.17 | 0.59 | -0.32 | 0.29 | -0.67 | 0.01 |
| R-IP | 0.37 | 0.21 | -0.43 | 0.14 | 0.37 | 0.21 | -0.74 | 0.00 |
| R-PCUN | -0.23 | 0.46 | -0.56 | 0.05 | -0.43 | 0.14 | -0.74 | 0.00 |
| R-CUN | 0.04 | 0.90 | -0.15 | 0.62 | 0.25 | 0.41 | -0.35 | 0.25 |
| R-PCA | 0.45 | 0.12 | 0.37 | 0.21 | 0.28 | 0.35 | -0.31 | 0.30 |
| R-LOCC | 0.24 | 0.43 | 0.32 | 0.28 | -0.07 | 0.83 | 0.03 | 0.93 |
| R-LING | 0.64 | 0.02 | 0.20 | 0.52 | 0.38 | 0.20 | -0.58 | 0.04 |
| R-FUS | 0.16 | 0.59 | -0.51 | 0.08 | -0.13 | 0.68 | 0.30 | 0.33 |
| R-PARH | -0.51 | 0.08 | -0.52 | 0.07 | -0.58 | 0.04 | -0.33 | 0.27 |
| R-ENT | -0.46 | 0.11 | -0.39 | 0.19 | -0.47 | 0.11 | 0.61 | 0.03 |
| R-TP | -0.03 | 0.93 | -0.33 | 0.28 | -0.18 | 0.56 | -0.29 | 0.34 |
| R-IT | -0.68 | 0.01 | -0.45 | 0.12 | -0.57 | 0.04 | -0.30 | 0.32 |
| R-MT | -0.13 | 0.68 | -0.53 | 0.06 | -0.27 | 0.38 | -0.76 | 0.00 |
| R-BSTS | 0.29 | 0.34 | -0.08 | 0.79 | -0.07 | 0.81 | -0.19 | 0.53 |
| R-ST | 0.52 | 0.07 | -0.36 | 0.22 | 0.00 | 1.00 | -0.15 | 0.63 |
| R-TT | 0.00 | 0.99 | 0.07 | 0.83 | 0.17 | 0.59 | 0.08 | 0.79 |
| R-INS | 0.13 | 0.67 | -0.01 | 0.97 | -0.11 | 0.72 | -0.26 | 0.40 |
| R-THA | -0.31 | 0.30 | -0.09 | 0.76 | -0.53 | 0.06 | 0.52 | 0.07 |
| R-CAU | -0.40 | 0.17 | -0.16 | 0.60 | -0.36 | 0.22 | -0.48 | 0.10 |
| R-PUT | -0.67 | 0.01 | -0.40 | 0.18 | -0.28 | 0.36 | 0.33 | 0.26 |
| R-PAL | -0.50 | 0.08 | -0.71 | 0.01 | -0.50 | 0.08 | 0.03 | 0.93 |
| R-ACCUM | -0.74 | 0.00 | -0.48 | 0.10 | -0.56 | 0.05 | -0.16 | 0.60 |
| R-HIP | 0.33 | 0.27 | -0.59 | 0.03 | -0.27 | 0.38 | -0.18 | 0.57 |
| R-AMYG | -0.20 | 0.51 | -0.41 | 0.17 | -0.57 | 0.04 | -0.13 | 0.68 |
| L-LOF | 0.33 | 0.27 | -0.29 | 0.33 | -0.20 | 0.51 | -0.29 | 0.34 |
| L-PORB | 0.32 | 0.29 | -0.23 | 0.44 | -0.19 | 0.52 | -0.37 | 0.21 |
| L-FP | 0.28 | 0.36 | 0.38 | 0.20 | -0.16 | 0.61 | 0.01 | 0.97 |
| L-MOF | 0.14 | 0.66 | -0.51 | 0.08 | -0.35 | 0.25 | -0.19 | 0.54 |
| L-PTRI | -0.15 | 0.62 | -0.61 | 0.03 | -0.20 | 0.52 | -0.64 | 0.02 |
| L-POPE | 0.21 | 0.49 | -0.30 | 0.32 | -0.08 | 0.80 | -0.37 | 0.21 |
| L-RMF | 0.16 | 0.61 | -0.22 | 0.47 | -0.42 | 0.16 | -0.79 | 0.00 |
| L-SF | 0.40 | 0.18 | -0.72 | 0.01 | -0.01 | 0.97 | -0.33 | 0.27 |
| L-CMF | 0.59 | 0.03 | -0.43 | 0.14 | 0.50 | 0.08 | -0.38 | 0.20 |
| L-PREC | 0.30 | 0.31 | 0.06 | 0.86 | -0.35 | 0.25 | -0.58 | 0.04 |
| L-PARC | -0.30 | 0.31 | 0.04 | 0.89 | -0.37 | 0.22 | 0.13 | 0.68 |
| L-RAC | -0.51 | 0.08 | -0.41 | 0.16 | -0.47 | 0.11 | -0.13 | 0.67 |
| L-CAC | -0.25 | 0.41 | -0.32 | 0.29 | -0.19 | 0.53 | -0.27 | 0.38 |
| L-PC | -0.19 | 0.54 | -0.13 | 0.67 | -0.45 | 0.13 | -0.28 | 0.35 |
| L-ISTC | 0.02 | 0.94 | -0.32 | 0.29 | 0.04 | 0.89 | -0.51 | 0.07 |
| L-PSTC | 0.42 | 0.15 | 0.57 | 0.04 | 0.12 | 0.69 | 0.44 | 0.14 |
| L-SMAR | -0.18 | 0.57 | -0.66 | 0.01 | -0.15 | 0.61 | -0.62 | 0.02 |
| L-SP | 0.26 | 0.38 | 0.37 | 0.21 | -0.28 | 0.36 | -0.34 | 0.25 |
| L-IP | 0.03 | 0.93 | -0.54 | 0.06 | -0.01 | 0.98 | -0.62 | 0.02 |
| L-PCUN | 0.12 | 0.69 | -0.46 | 0.12 | 0.02 | 0.96 | -0.19 | 0.54 |
| L-CUN | 0.17 | 0.58 | -0.03 | 0.92 | 0.26 | 0.39 | -0.64 | 0.02 |
| L-PCA | 0.34 | 0.26 | 0.26 | 0.38 | -0.15 | 0.63 | -0.23 | 0.46 |
| L-LOCC | 0.10 | 0.74 | 0.19 | 0.53 | -0.45 | 0.12 | -0.21 | 0.50 |
| L-LING | 0.45 | 0.12 | 0.29 | 0.34 | -0.03 | 0.92 | 0.09 | 0.77 |
| L-FUS | -0.19 | 0.54 | -0.58 | 0.04 | -0.40 | 0.18 | -0.46 | 0.11 |
| L-PARH | -0.02 | 0.95 | -0.44 | 0.13 | -0.22 | 0.47 | -0.31 | 0.30 |
| L-ENT | 0.07 | 0.82 | -0.40 | 0.18 | -0.10 | 0.74 | -0.07 | 0.81 |
| L-TP | -0.08 | 0.79 | -0.65 | 0.02 | -0.04 | 0.89 | -0.46 | 0.12 |
| L-IT | -0.48 | 0.10 | -0.50 | 0.08 | -0.47 | 0.10 | -0.23 | 0.45 |
| L-MT | 0.05 | 0.86 | -0.09 | 0.78 | -0.35 | 0.24 | -0.04 | 0.91 |
| L-BSTS | 0.05 | 0.88 | 0.09 | 0.77 | -0.32 | 0.29 | -0.17 | 0.58 |
| L-ST | 0.42 | 0.15 | -0.62 | 0.02 | 0.16 | 0.60 | -0.60 | 0.03 |
| L-TT | -0.05 | 0.86 | 0.23 | 0.45 | 0.34 | 0.26 | 0.02 | 0.95 |
| L-INS | 0.00 | 0.99 | -0.47 | 0.10 | -0.11 | 0.72 | -0.18 | 0.56 |
| L-THA | -0.30 | 0.32 | -0.28 | 0.36 | -0.49 | 0.09 | 0.59 | 0.03 |
| L-CAU | -0.60 | 0.03 | -0.31 | 0.31 | -0.36 | 0.22 | 0.32 | 0.29 |
| L-PUT | -0.30 | 0.32 | -0.37 | 0.22 | -0.44 | 0.13 | 0.14 | 0.64 |
| L-PAL | -0.86 | 0.00 | 0.28 | 0.35 | -0.71 | 0.01 | 0.67 | 0.01 |
| L-ACCUM | 0.09 | 0.77 | -0.37 | 0.21 | -0.23 | 0.46 | 0.04 | 0.89 |
| L-HIP | 0.28 | 0.35 | -0.24 | 0.42 | -0.09 | 0.77 | 0.13 | 0.68 |
| L-AMYG | 0.29 | 0.34 | 0.23 | 0.46 | 0.25 | 0.41 | -0.14 | 0.64 |

Supplementary Table 3 - Group comparisons between bipolar disorder patients and healthy controls (controlled for age, uncorrected p-values)

| regions | Structural connectivity |  | Functional Connectivity |  | EC input-strength |  | EC output-strength |  | Node variance |  |
| --- | --- | --- | --- | --- | --- | --- | --- | --- | --- | --- |
|  | T-val | p-val | T-val | p-val | T-val | p-val | T-val | p-val | T-val | p-val |
| R-LOF | 2.06 | 0.06 | 1.64 | 0.11 | 0.01 | 0.99 | -0.24 | 0.82 | -2.34 | 0.02 |
| R-PORB | 0.44 | 0.69 | 1.70 | 0.11 | -0.86 | 0.39 | 1.30 | 0.20 | -2.34 | 0.03 |
| R-FP | 0.96 | 0.38 | 1.41 | 0.17 | 0.97 | 0.33 | 0.16 | 0.87 | -0.99 | 0.35 |
| R-MOF | 0.08 | 0.94 | 1.37 | 0.19 | 0.40 | 0.71 | 0.18 | 0.87 | -1.94 | 0.06 |
| R-PTRI | 0.67 | 0.52 | 1.89 | 0.07 | 1.60 | 0.13 | 1.25 | 0.23 | -2.26 | 0.02 |
| R-POPE | -0.49 | 0.62 | 1.53 | 0.14 | 0.09 | 0.93 | -0.01 | 0.99 | -1.84 | 0.07 |
| R-RMF | 1.57 | 0.13 | 1.39 | 0.18 | 0.53 | 0.60 | -1.03 | 0.33 | -1.71 | 0.11 |
| R-SF | 1.78 | 0.10 | 1.18 | 0.25 | 0.42 | 0.70 | -0.87 | 0.40 | -2.08 | 0.04 |
| R-CMF | 0.92 | 0.38 | 1.39 | 0.18 | 0.00 | 1.00 | 0.36 | 0.73 | -1.03 | 0.33 |
| R-PREC | 0.60 | 0.56 | 1.66 | 0.11 | 0.43 | 0.66 | -1.54 | 0.14 | -1.54 | 0.14 |
| R-PARC | 0.61 | 0.55 | 0.99 | 0.33 | -0.54 | 0.60 | -0.27 | 0.80 | -1.17 | 0.26 |
| R-RAC | 1.94 | 0.07 | 1.47 | 0.15 | 0.01 | 0.99 | 0.27 | 0.78 | -2.08 | 0.04 |
| R-CAC | 0.58 | 0.57 | 0.99 | 0.33 | 0.41 | 0.68 | 0.51 | 0.61 | -2.08 | 0.04 |
| R-PC | 0.73 | 0.47 | 1.88 | 0.08 | -1.41 | 0.17 | 0.64 | 0.53 | -1.62 | 0.09 |
| R-ISTC | 0.26 | 0.79 | 1.21 | 0.25 | -0.85 | 0.42 | 0.53 | 0.62 | -1.48 | 0.15 |
| R-PSTC | 1.30 | 0.21 | 1.62 | 0.12 | 0.92 | 0.37 | -1.15 | 0.26 | -0.82 | 0.42 |
| R-SMAR | -0.19 | 0.84 | 1.30 | 0.21 | -0.56 | 0.58 | -0.36 | 0.73 | -1.52 | 0.15 |
| R-SP | -0.12 | 0.91 | 1.31 | 0.20 | 0.43 | 0.69 | -1.67 | 0.12 | -1.03 | 0.38 |
| R-IP | 0.20 | 0.85 | 1.44 | 0.16 | 0.18 | 0.87 | -0.85 | 0.43 | -1.24 | 0.23 |
| R-PCUN | 1.72 | 0.11 | 1.97 | 0.06 | -0.73 | 0.47 | -0.39 | 0.72 | -1.57 | 0.11 |
| R-CUN | 1.48 | 0.15 | 2.09 | 0.05 | -0.74 | 0.46 | -1.07 | 0.30 | -1.21 | 0.24 |
| R-PCA | 1.28 | 0.21 | 2.39 | 0.03 | -0.90 | 0.37 | -0.30 | 0.76 | -1.59 | 0.11 |
| R-LOCC | -0.12 | 0.91 | 1.43 | 0.18 | 0.75 | 0.47 | -1.51 | 0.14 | -1.00 | 0.38 |
| R-LING | 0.94 | 0.37 | 1.89 | 0.08 | -0.35 | 0.74 | -0.76 | 0.46 | -1.17 | 0.28 |
| R-FUS | 0.28 | 0.78 | 1.38 | 0.19 | -1.69 | 0.11 | 0.19 | 0.85 | -2.63 | 0.01 |
| R-PARH | 0.88 | 0.39 | 0.79 | 0.45 | -0.37 | 0.72 | 0.10 | 0.91 | -1.76 | 0.06 |
| R-ENT | -0.77 | 0.47 | 0.28 | 0.78 | -1.05 | 0.31 | 0.11 | 0.92 | -1.76 | 0.10 |
| R-TP | -1.75 | 0.09 | 1.39 | 0.17 | -1.39 | 0.18 | 0.20 | 0.85 | -1.89 | 0.07 |
| R-IT | -0.13 | 0.90 | 1.25 | 0.23 | -1.49 | 0.16 | 0.67 | 0.50 | -2.66 | 0.01 |
| R-MT | 1.00 | 0.34 | 2.16 | 0.04 | -1.02 | 0.33 | -0.02 | 0.98 | -2.41 | 0.02 |
| R-BSTS | -0.31 | 0.76 | 1.98 | 0.06 | 0.70 | 0.50 | 0.91 | 0.38 | -1.32 | 0.20 |
| R-ST | 0.83 | 0.41 | 2.03 | 0.06 | -0.31 | 0.77 | -0.14 | 0.90 | -1.64 | 0.11 |
| R-TT | 0.29 | 0.77 | 2.39 | 0.03 | -0.32 | 0.75 | 1.59 | 0.13 | -2.08 | 0.04 |
| R-INS | 0.81 | 0.43 | 1.78 | 0.09 | -0.78 | 0.44 | 0.34 | 0.73 | -1.50 | 0.15 |
| R-THA | 0.60 | 0.54 | 0.84 | 0.41 | -0.26 | 0.81 | 0.42 | 0.68 | -1.62 | 0.09 |
| R-CAU | 1.50 | 0.14 | 0.58 | 0.56 | -0.48 | 0.63 | 0.01 | 0.99 | -2.18 | 0.03 |
| R-PUT | 0.85 | 0.40 | 0.67 | 0.50 | -1.01 | 0.35 | -1.56 | 0.13 | -1.68 | 0.08 |
| R-PAL | -1.19 | 0.25 | 0.09 | 0.93 | -1.15 | 0.26 | -0.36 | 0.73 | -1.84 | 0.06 |
| R-ACCUM | 0.17 | 0.86 | 0.49 | 0.62 | 0.54 | 0.58 | 1.66 | 0.11 | -2.08 | 0.05 |
| R-HIP | 0.19 | 0.85 | 0.72 | 0.48 | -0.33 | 0.75 | -0.72 | 0.49 | -2.07 | 0.04 |
| R-AMYG | -0.42 | 0.69 | 1.66 | 0.11 | -1.47 | 0.16 | 0.32 | 0.75 | -2.25 | 0.03 |
| L-LOF | 0.61 | 0.55 | 1.68 | 0.11 | 0.09 | 0.93 | -0.41 | 0.69 | -1.97 | 0.05 |
| L-PORB | 1.98 | 0.06 | 2.46 | 0.02 | 0.35 | 0.73 | 1.20 | 0.24 | -2.21 | 0.03 |
| L-FP | 1.88 | 0.07 | 1.20 | 0.24 | 0.67 | 0.51 | -0.28 | 0.77 | -0.62 | 0.56 |
| L-MOF | 1.72 | 0.11 | 1.70 | 0.10 | -0.56 | 0.57 | 0.13 | 0.91 | -1.71 | 0.09 |
| L-PTRI | 1.97 | 0.06 | 1.74 | 0.10 | 0.88 | 0.38 | 0.23 | 0.83 | -1.92 | 0.07 |
| L-POPE | 0.41 | 0.71 | 1.39 | 0.18 | -0.48 | 0.64 | -0.55 | 0.61 | -1.76 | 0.08 |
| L-RMF | 1.18 | 0.25 | 1.81 | 0.07 | 0.57 | 0.58 | 0.18 | 0.87 | -1.77 | 0.08 |
| L-SF | 2.03 | 0.05 | 1.19 | 0.24 | 0.06 | 0.96 | -1.01 | 0.33 | -2.19 | 0.04 |
| L-CMF | 0.07 | 0.94 | 1.35 | 0.20 | -0.24 | 0.81 | -0.35 | 0.72 | -1.30 | 0.23 |
| L-PREC | 0.85 | 0.40 | 1.84 | 0.08 | -0.33 | 0.76 | 0.14 | 0.89 | -2.22 | 0.03 |
| L-PARC | -0.93 | 0.37 | 0.93 | 0.37 | -0.36 | 0.72 | -0.81 | 0.43 | -1.49 | 0.13 |
| L-RAC | 1.73 | 0.10 | 1.12 | 0.28 | 0.09 | 0.93 | 0.45 | 0.66 | -2.08 | 0.05 |
| L-CAC | 0.66 | 0.52 | 1.25 | 0.22 | 0.22 | 0.83 | 0.93 | 0.36 | -1.91 | 0.07 |
| L-PC | 2.17 | 0.04 | 1.98 | 0.06 | -1.29 | 0.21 | 1.00 | 0.32 | -1.26 | 0.22 |
| L-ISTC | 2.41 | 0.02 | 1.78 | 0.09 | -0.44 | 0.67 | 0.84 | 0.40 | -1.70 | 0.10 |
| L-PSTC | 0.42 | 0.67 | 1.37 | 0.18 | 0.29 | 0.77 | -1.05 | 0.31 | -1.28 | 0.22 |
| L-SMAR | 1.21 | 0.23 | 1.23 | 0.24 | -0.31 | 0.76 | -0.60 | 0.55 | -1.81 | 0.08 |
| L-SP | 0.35 | 0.72 | 1.38 | 0.18 | -0.51 | 0.62 | -1.15 | 0.26 | -1.94 | 0.07 |
| L-IP | 0.28 | 0.79 | 1.54 | 0.14 | -0.18 | 0.86 | -0.58 | 0.57 | -1.17 | 0.26 |
| L-PCUN | -0.43 | 0.67 | 2.37 | 0.03 | -0.04 | 0.97 | -0.04 | 0.97 | -1.49 | 0.14 |
| L-CUN | 1.43 | 0.17 | 1.83 | 0.08 | -1.32 | 0.20 | -0.55 | 0.57 | -1.68 | 0.10 |
| L-PCA | 0.43 | 0.67 | 2.43 | 0.03 | -0.58 | 0.56 | 0.22 | 0.83 | -1.62 | 0.10 |
| L-LOCC | 1.43 | 0.17 | 1.82 | 0.09 | -0.13 | 0.89 | -0.71 | 0.50 | -1.67 | 0.12 |
| L-LING | -0.22 | 0.84 | 2.13 | 0.05 | 0.82 | 0.42 | -0.01 | 0.99 | -1.53 | 0.12 |
| L-FUS | 0.17 | 0.87 | 1.37 | 0.18 | -1.46 | 0.17 | -0.17 | 0.86 | -0.46 | 0.86 |
| L-PARH | 0.82 | 0.42 | 0.89 | 0.40 | 0.07 | 0.95 | 0.81 | 0.41 | -2.09 | 0.03 |
| L-ENT | -0.15 | 0.88 | 0.16 | 0.86 | -0.89 | 0.41 | -0.14 | 0.93 | -0.91 | 0.46 |
| L-TP | -0.66 | 0.52 | 0.41 | 0.69 | -0.72 | 0.52 | -0.91 | 0.36 | -1.11 | 0.31 |
| L-IT | -1.09 | 0.29 | 1.16 | 0.26 | -1.00 | 0.34 | 0.29 | 0.77 | -1.07 | 0.35 |
| L-MT | 1.16 | 0.26 | 1.97 | 0.06 | -0.22 | 0.84 | -0.03 | 0.98 | -2.16 | 0.04 |
| L-BSTS | -0.66 | 0.51 | 2.28 | 0.03 | -0.59 | 0.59 | 1.30 | 0.21 | -1.40 | 0.17 |
| L-ST | -0.14 | 0.90 | 2.28 | 0.04 | 0.74 | 0.47 | 0.63 | 0.55 | -2.04 | 0.06 |
| L-TT | -0.59 | 0.60 | 1.92 | 0.08 | 0.36 | 0.72 | 1.51 | 0.14 | -1.45 | 0.16 |
| L-INS | 0.95 | 0.35 | 1.34 | 0.20 | -1.12 | 0.28 | -0.65 | 0.55 | -2.35 | 0.01 |
| L-THA | 2.10 | 0.05 | 1.11 | 0.27 | -1.24 | 0.25 | 0.04 | 0.97 | -1.38 | 0.15 |
| L-CAU | 0.52 | 0.61 | 0.50 | 0.61 | -1.40 | 0.18 | -0.31 | 0.76 | -2.58 | 0.01 |
| L-PUT | 0.37 | 0.72 | 1.43 | 0.17 | -1.42 | 0.18 | -0.23 | 0.82 | -2.39 | 0.02 |
| L-PAL | -1.94 | 0.06 | -0.22 | 0.83 | -0.78 | 0.50 | -0.67 | 0.49 | -1.36 | 0.17 |
| L-ACCUM | 0.05 | 0.95 | 0.53 | 0.59 | -0.36 | 0.73 | 0.37 | 0.70 | -1.39 | 0.17 |
| L-HIP | 1.22 | 0.26 | 0.52 | 0.62 | -0.04 | 0.96 | -0.64 | 0.53 | -1.75 | 0.09 |
| L-AMYG | 0.58 | 0.84 | 0.83 | 0.42 | -1.13 | 0.26 | -0.04 | 0.96 | -2.53 | 0.01 |

## References

Benjamini, Yoav, and Daniel Yekutieli, 2001. “The Control of the False Discovery Rate in Multiple Testing under Dependency.” The Annals of Statistics. 29 (4) 1165–1188 doi:10.1214/aos/1013699998

Biswal, B., F.Z. Yetkin, V.M. Haughton, and J.S. Hyde, 1995. “Functional Connectivity in the Motor Cortex of Resting Human Brain Using Echo-Planar MRI.” Magnetic Resonance in Medicine: Official Journal of the Society of Magnetic Resonance in Medicine/Society of Magnetic Resonance in Medicine. 34 (4) 537–41

Julio, Bobes, Antonio Bulbena, Antonio Luque, Rafael Dal-Ré, Javier Ballesteros, Nora Ibarra, and Grupo De Validacion En Espanol De Escalas Psicometricas, 2003. “[A comparative psychometric study of the Spanish versions with 6, 17, and 21 items of the Hamilton Depression Rating Scale].” Medicina Clfnica. 120 (18) 693–700

Bright, G.Molly, Marta Bianciardi, Jacco A. de Zwart, Kevin Murphy, and H. Duyn Jeff, 2014. “Early Anti-Correlated BOLD Signal Changes of Physiologic Origin.” Neuroimage. 87 February 287–96 doi:10.1016/j.neuroimage.2013.10.055

R.B. Buxton, E.C. Wong, and L.R. Frank, 1998. “Dynamics of Blood Flow and Oxygenation Changes during Brain Activation: The Balloon Model.” Magnetic Resonance in Medicine. 39 (6) 855–64 doi:10.1002/mrm.1910390602

Cabral, Joana, L. Morten, Kringelbach, and Gustavo Deco, 2014. “Exploring the Network Dynamics Underlying Brain Activity during Rest.” Progress in Neurobiology. 114 March 102–131 doi:10.1016/j.pneurobio.2013.12.005

Chang, Catie, and H. Glover Gary, 2010. “Time-frequency Dynamics of Resting-State Brain Connectivity Measured with fMRI.” Neuroimage. 50 (1) 81–98 doi:10.1016/j.neuroimage.2009.12.011

Colom, Francesc, Eduard Vieta, Anabel Martímez-Arán, Margarida Garcia-Garcia, Manía Reinares, Carla Torrent, José Manuel Goikolea, Sebastià Banús, and Manel Salamero, 2002. “[Spanish version of a scale for the assessment of mania: validity and reliability of the Young Mania Rating Scale].” Medicina Clfnica. 119 (10) 366–71

E. Damaraju, E.A. Allen, A. Belger, J.M. Ford, S. McEwen, D.H. Mathalon, B.A. Mueller, et al.2014. “Dynamic Functional Connectivity Analysis Reveals Transient States of Dysconnectivity in Schizophrenia.” Neuroimage: Clinical. (5) 298–308 doi:10.1016/J.Nicl.2014.07.003

Deco, Gustavo, and Viktor K. Jirsa, 2012. “Ongoing Cortical Activity at Rest: Criticality, Multistability, and Ghost Attractors.” The Journal of Neuroscience. 32 (10) 3366–75 doi:10.1523/JNEUROSCI.2523-11.2012

Deco, Gustavo, Viktor Jirsa, A.R. McIntosh, and Olaf Sporns, Rölf Kotter, 2009. “Key Role of Coupling, Delay, and Noise in Resting Brain Fluctuations.” Proceedings of the National Academy of Sciences. 106 (25) 10302–7 doi:10.1073/pnas.0901831106

Deco, Gustavo, Anthony R McIntosh, Kelly Shen, R. Matthew Hutchison, Ravi S. Menon, Stefan Everling, Patric Hagmann, and Viktor K. Jirsa, 2014. “Identification of Optimal Structural Connectivity Using Functional Connectivity and Neural Modeling.” The Journal of Neuroscience. 34 (23) 7910–16 doi:10.1523/JNEUROSCI.4423-13.2014

Demirtaş, Murat, Cristian Tornador, Carles Falcón, Marina López-Solà, Rosa Hernández-Ribas, Jesús Pujol, José M. Menchón, et al.2016. “Dynamic Functional Connectivity Reveals Altered Variability in Functional Connectivity among Patients with Major Depressive Disorder.” Human Brain Mapping. 37 (8) 2918–30 doi:10.1002/Hbm.23215

M.B. First, 2001.“Structured Clinical Interview for DSM-IV-TR Axis Disorders, Research Version,Patient Edition (SCID-I/P).”. New York: New York State Psychiatric Institute

M.B. First, 2002.“Structured Clinical Interview for DSM-IV-TR Axis Disorders, Research Version,Patient Edition (SCID-I/P).”. New York: New York State Psychiatric Institute

Fox, D. Michael, and Michael Greicius, 2010. “Clinical Applications of Resting State Functional Connectivity.” Frontiers in Systems Neuroscience. 419 doi:10.3389/fnsys.2010.00019

Fox, D. Michael, and Marcus E. Raichle, 2007. “Spontaneous Fluctuations in Brain Activity Observed with Functional Magnetic Resonance Imaging.” Nature Reviews Neuroscience. 8 (9) 700–711 doi:10.1038/nrn2201

Fox, D. Michael, Abraham Z. Snyder, Justin L. Vincent, Maurizio Corbetta, David C. Van Essen, and Marcus E. Raichle, 2005. “The Human Brain Is Intrinsically Organized into Dynamic, Anticorrelated Functional Networks.” Proceedings of the National Academy of Sciences of the United States of America. 102 (27) 9673–78 doi:10.1073/pnas.0504136102

Fox, D. Michael, Dongyang Zhang, Abraham Z. Snyder, Marcus E. Raichle, 2009. “The Global Signal and Observed Anticorrelated Resting State Brain Networks.” Journal of Neurophysiology. 101 (6) 3270–83 doi:10.1152/jn.90777.2008

Friston, J. Karl, 2011. “Functional and Effective Connectivity: A Review.” Brain Connectivity. 1 (1) 13–36 doi:10.1089/brain.2011.0008

Friston, J. Karl, Joshua Kahan, Bharat Biswal, and Adeel Razi, 2014. “A DCM for Resting State fMRI.” Neuroimage. 94 July396–407 doi:10.1016/j.neuroimage.2013.12.009.

K.J. Friston, L. Harrison, and W. Penny, 2003. “Dynamic Causal Modelling.” Neuroimage. 19 (4) 1273–1302

Ghosh, Anandamohan, Y. Rho, A.R. McIntosh, R. Kötter, and V.K. Jirsa, 2008. “Noise during Rest Enables the Exploration of the Brain’s Dynamic Repertoire.” Plos Comput Biol. 4 (10) E1000196 doi:10.1371/Journal.Pcbi.1000196

Gilson, Matthieu, Ruben Moreno-Bote, Adrián Ponce-Alvarez, Petra Ritter, and Gustavo Deco, 2016. “Estimation of Directed Effective Connectivity from fMRI Functional Connectivity Hints at Asymmetries of Cortical Connectome.” PLOS Comput Biol 12 (3): E1004762. Doi:10.1371/Journal.Pcbi.1004762. 12 (3) e1004762 doi:10.1371/journal.pcbi.1004762.

Gopinath, Kaundinya, Venkatagiri Krishnamurthy, Romeo Cabanban, and Bruce A. Crosson, 2015. “Hubs of Anticorrelation in High-Resolution Resting-State Functional Connectivity Network Architecture.” Brain Connectivity. 5 (5) 267–75 doi:10.1089/brain.2014.0323

Greicius, Michael, 2008. “Resting-State Functional Connectivity in Neuropsychiatric Disorders.” Current Opinion in Neurology. 21 (4) 424–30 doi:10.1097/WCO.0b013e328306f2c5

Hansen, C.A. Enrique, Demian Battaglia, Reas Spiegler, Gustavo Deco, and Viktor K. Jirsa, 2015. “Functional Connectivity Dynamics: Modeling the Switching Behavior of the Resting State.” Neuroimage. 105 January 525–35 doi:10.1016/j.neuroimage.2014.11.001

Honey, J. Christopher, Rolf Kötter, Michael Breakspear, and Olaf Sporns, 2007. “Network Structure of Cerebral Cortex Shapes Functional Connectivity on Multiple Time Scales.” Proceedings of the National Academy of Sciences. 104 (24) 10240–45 doi:10.1073/pnas.0701519104

C.J. Honey, O. Sporns, L. Cammoun, X. Gigandet, J.P. Thiran, and R. Meuli, P. Hagmann, 2009. “Predicting Human Resting-State Functional Connectivity from Structural Connectivity.” Proceedings of the National Academy of Sciences. 106 (6) 2035–40 doi:10.1073/pnas.0811168106

Hutchison, R. Matthew, Thilo Womelsdorf, Joseph S. Gati, Stefan Everling, and Ravi S. Menon, 2013. “Resting-State Networks Show Dynamic Functional Connectivity in Awake Humans and Anesthetized Macaques.” Human Brain Mapping. 34 (9) 2154–77 doi:10.1002/Hbm.22058

N.T. Markov, M.M. Ercsey-Ravasz, A.R. Ribeiro Gomes, C. Lamy, L. Magrou, J. Vezoli, P. Misery, 2014.et al.“A Weighted and Directed Interareal Connectivity Matrix for Macaque Cerebral Cortex.” Cerebral Cortex. 24 (1) 17–36 doi:10.1093/cercor/bhs270

Messé, Arnaud, Habib Benali, and Guillaume Marrelec, “Relating Structural and Functional Connectivity in MRI: A Simple Model for a Complex Brain.” IEEE Transactions on Medical Imaging. 2015.34 (1) 27–37 doi:10.1109/TMI.2014.2341732

Murphy, Kevin, Rasmus M. Birn, Daniel A. Handwerker, Tyler B. Jones, and Peter A. Bandettini, 2009. “The Impact of Global Signal Regression on Resting State Correlations: Are Anti-Correlated Networks Introduced?”Neuroimage. 44 (3) 893–905 doi:10.1016/j.neuroimage.2008.09.036

F. Nikolaou, C. Orphanidou, P. Papakyriakou, K. Murphy, R.G. Wise, and G.D. Mitsis, 2016. “Spontaneous Physiological Variability Modulates Dynamic Functional Connectivity in Resting-State Functional Magnetic Resonance Imaging.” Phil.Trans. R. Soc. A. 374 (2067) 20150183 doi:10.1098/Rsta.2015.0183

D. Popovic, C. Torrent, J.M. Goikolea, N. Cruz, J. Sánchez-Moreno, A. González-Pinto, and E. Vieta, 2014. “Clinical Implications of Predominant Polarity and the Polarity Index in Bipolar Disorder: A Naturalistic Study.” Acta Psychiatrica Scandinavica. 129 (5) 366–74 doi:10.1111/Acps.12179

Power, Jonathan D, Mark Plitt, Timothy O Laumann, and Alex Martin, 2016. “Sources and Implications of Whole-Brain fMRI Signals in Humans.” Neuroimage. Accessed October. 28 doi:10.1016/J.Neuroimage.2016.09.038

Prčkovska, Vesna, Paulo Rodrigues, Ana Puigdellivol Sanchez, Marc Ramos, Magi Andorra, Eloy Martinez-Heras, Carles Falcon, Albert Prats-Galino, and Pablo Villoslada, 2016. “Reproducibility of the Structural Connectome Reconstruction across Diffusion Methods.” Journal of Neuroimaging: Official Journal of the American Society of Neuroimaging. 26 (1) 46–57 doi:10.1111/jon.12298

Rosa, R. Adriane, Jose Sánchez-Moreno, Anabel Martĺmez-Aran, Manel Salamero, Carla Torrent, Maria Reinares, Mercé Comes, et al.2007. “Validity and Reliability of the Functioning Assessment Short Test (FAST) in Bipolar Disorder.” Clinical Practice and Epidemiology in Mental Health: CP & EMH3 (June). 5 doi:10.1186/1745-0179-3-5

Uddin, Q. Lucina, A.M. Clare Kelly, Bharat B. Biswal, and F. Xavier Castellanos, Michael P. Milham, 2009. “Functional Connectivity of Default Mode Network Components: Correlation, Anticorrelation, and Causality.” Human Brain Mapping. 30 (2) 625–37 doi:10.1002/Hbm.20531

Van Essen, C. David, Saad Jbabdi, Stamatios N. Sotiropoulos, Charles Chen, Krikor Dikranian, Tim Coalson, John Harwell, Timothy E. J. Behrens, and Matthew F. Glasser, 2014. “Mapping Connections in Humans and NonHuman Primates: Aspirations and Challenges for Diffusion Imaging.” Diffusion MRI (Second Edition). 337–58http://www.sciencedirect.com/science/article/pii/B9780123964601000160

E. Vieta Pascual, C. Torrent Font, A. Martmez-Aran, F. Colom Victoriano, and M. Reinares Gabnepen, A. Benabarre Hernández, M. Comes Forastero, J.M. Goikolea Alberdi, 2002. “[A user-friendly scale for the short and long term outcome of bipolar disorder: the CGI-BP-M].” Actas Espñnolas De Psiquiatria. 30 (5) 301–4

Whitfield-Gabrieli, Susan, and Judith M. Ford, “Default Mode Network Activity and Connectivity in Psychopathology.” Annual Review of Clinical Psychology. 2012.8 (1) 49–76 doi:10.1146/annurev-clinpsy-032511-143049

Yang, J. Genevieve, John D. Murray, Grega Repovs, Michael W. Cole, Aleksandar Savic, Matthew F. Glasser, Christopher Pittenger, et al.2014. “Altered Global Brain Signal in Schizophrenia.” Proceedings of the National Academy of Sciences of the United States of America. 111 (20) 7438–43 doi:10.1073/Pnas.1405289111

Zalesky, Andrew, Alex Fornito, Luca Cocchi, Leonardo L. Gollo, and Michael Breakspear, 2014. “Time-Resolved Resting-State Brain Networks.” Proceedings of the National Academy of Sciences. 111 (28) 10341–46 doi:10.1073/Pnas.1400181111

